# Are bio-sourced nanoplastics inert for aquatic species? A toxicity study on three micro-algae species and a freshwater bivalve

**DOI:** 10.64898/2026.06.14.732166

**Authors:** Adeline Arini, Anderson M. Medeiros, Véronique Coma, Etienne Grau, Olivier Sandre, Magalie Baudrimont

## Abstract

Concerns raised by ubiquitate plastic contamination are urging to develop alternative materials. In the recent years, bio-sourced polymers also coined as “bioplastics” have been proposed to mitigate plastic pollution while meeting industrial and commercial expectations. Like petro-sourced plastics, they are expected to break-down in the environment into fragments down to sub-micron size. However, only scarce data are available on the impacts of such biosourced nanoplastics once released into the environment. This study examines the effects on aquatic species of model nanoplastics made from several bio-sourced polymers (Bio-NPs) that are either already on market (PHA, PLA, PA11) or still under development (NIPU, PCAR). We exposed three species of micro-algae (at 10, 100, and 1000 µg/L, for 24 and 48 hours, and one week) to test the effects of Bio-NPs on algal growth. We also exposed freshwater bivalves *C. fluminea* (at 1, 10 and 100 µg/L, for one week) to test the filtration activity and gene expressions in response to Bio-NPs exposure. All five Bio-NPs tested generated growth inhibitions in at least one of the three algae tested. PLA and PA11 were the most deleterious ones for algal growth among the five tested Bio-NPs. The highest growth inhibitions were observed on the fresh water species *D. subspicatus*. Each Bio-NP tested resulted in significant decreases of the filtration rates of *C. fluminea*. PHA impaired filtration at the lowest concentrations tested (1 µg/L) whereas PCAR, PA11 and NIPU led to significant effects only at higher concentrations (10 and 100 µg/L).

The results from gene expressions in *C. fluminea* showed strong inductions of all gene functions tested for all the five bio-NPs tested. These Bio-NPs triggered endocytosis and detoxification mechanisms. They impaired the mitochondrial metabolism and triggered oxidative stress and immune responses. PA11, NIPU and PHA exposures resulted in the strongest gene regulations. The present study brings brand new findings about a kind of nanoplastics that may be released into the environment in a near future as the use of bioplastics is growing fast. It will help better understanding the impacts of such fragmented bioplastic NPs on aquatic species.

## INTRODUCTION

Research is currently developing in the plastics sector to offer a new range of plastics made from renewable sources (plants, wood lignocellulosic derivatives, vegetable oils, etc.). Unlike petro-sourced plastics, derived from fossil materials, bio-sourced plastics are compounds synthesized from totally or partially renewable molecular bricks (Shah et al., 2021). Among materials called “bioplastics”, only a part of them are supposedly entirely biodegradable polymers that can be degraded by the enzymatic activity of microorganisms (Tournier et al., 2023). Altough some bioplastics with some hydrolysable bonds can effectively undergo biodegradation in a reasonable time under given conditions, most of them actually present similar physical and chemical properties as petroleum-based plastics. As such, bioplastics often regarded as valuable alternatives for petrosourced materials may be disappointing to really lower the plastic footprint (Mascarenhas & Aruna, 2017). It is important to mention that herein we only consider new polymers from biomass and exclude from the stuly “drop-in” bio-based plastic such as bio-polyethylene which is expected to present the same environmental impact as the petroleum polyethylene.

The main bioplastics available to date on market are polyesters, namely polyhydroxyalcanoates (PHA), polybutylene succinate (PBS) and poly(lactic acid) (PLA). They can be blended with natural starch, to obtain “starch plastics” or “starch blends”. Among those, PLA remains the most commercially mature bioplastic (European_bioplastics, 2023). Bioplastics exhibit innovative structures, compared to existing petrochemical polymers, while presenting similar performances for technical applications (European_bioplastics, 2023). Their greatest interests are to counteract CO2 emissions, the depletion of oil resources and seemingly to preserve the environment. Bioplastics are used for a growing variety of applications (packaging, electronics, automotive and textiles). Yet, they still currently represent less than one percent of total global thermoplastic production, which amounts to more than 390 million tons per year (European_bioplastics, 2023). However, the market for bioplastics continues to increase and forecasts predict an increasing production of 20 to 25% per year in the future years (Shah et al., 2021).

Plastic wastes are nowadays greatly known to degrade and break down into smaller pieces, reaching sub-micronic scale (Gigault et al., 2018). Despite a growing number of publications on petroleum-based nanoplastics (NPs) in recent years, many questions remain unanswered, not only in terms of toxicity, but also about their fate in the environment and the preventive measures to adopt to limit their diffusion into the environmental compartments (in water, in soil and the air). Due to the recent gain of interest for bioplastics, their fragmentation into NPs is not yet sufficiently considered by the environmental research community, and almost no data is available so far about their fate and potential toxicity once released into the environment. To date, reports on the impact of nanoplastics derived from bioplastics are scarce. The first published study reported damages caused by nanoplastics originating from one type of promising bio-sourced polyester (polyhydroxybutyrate (PHB)) to algal growth and inhibition of locomotion in the crustacean *Daphnia magna*, associated to oxidative stress (González-Pleiter et al., 2019). Another work studied the effets on planktons of leachates by microplastic suspensions of two common bioplastics, PLA and poly(hydroxybutyrate-*co*-valerate) (PHBV) together with polypropylene with a small percentage of nanoplastics (Laranjeiro et al., 2024) but the sub-micron particle fractions were minute, respectively 4%, 0.4% and 1.1% for PHBV, PLA, and PP. More recently, PLA nanoplastics were found not causing acute toxicity to marine rotifers, brine shrimps and zebrafish embryos (Mustapha et al., 2025) contrary to a previous study showing toxicity of PLA nanoplastics very close to those made of PP or LDPE on the locomotory activity of *Danio rerio* at concentrations as low as 1 µg/L at 120 hpf (Tamayo-Belda et al., 2023). Nonetheless, potential harmful effects of other biosourced nanoplastics remain unexplored to date, and especially on microalgae and bivalves. It is therefore urgent to carry out toxicity tests assessing the potential toxicity of bio-sourced NPs, before their large-scale production and marketing.

While ecotoxicological endpoints are necessary to establish their overall environmental impact, nanoplastics are still difficult to test due to absence of universal reference materials for them (Ducoli et al., 2025). In the literature reported so far (Crosset-Perrotin et al., 2025), there are two main route towards obtention of model nanoplastics for eco-toxicological studies: i) The top-down route where NPs are produced by fragmentation of virgin or aged plastics (e.g. litter collected from seashore or riverbanks) and successive sieving, fractionation or fitration steps until sub-µm size range; ii) Different bottom-up approaches to prepare NPs in-lab either by dispersed state polymerization (e.g. in emulsion, with surfactants) or by self-assembly of pre-synthesized polymer chains using the “nanoprecipitation” route, which consists in adding water as a bad solvent to a polymer solution (in a solvent miscible with water). In a recent work by Gálvez-Blanca et al (Gálvez-Blanca et al., 2026), PLA pellets were first cryo-grounded and sieved down to micropowder state, then they were photo-aged under UV light to produce PLA- NPs, which exhibited toxicity above 5 mg/L on *Pseudomas putida* bacteria. In the present study, both descending and ascending methods were combined to prepare a library of several bio-sourced NPs, either marketed or non-marketed. To the authors’ knowledge, it is the first report on the hazardous effects of a library of biosourced nanoplastics on two biological models: on the one hand *Corbicula fluminea*, a freshwater filter-feeding bivalve that possesses high accumulation capabilities, thus representing a suitable model used in many ecotoxicological studies to assess xenobiotic effects (Arini et al., 2020; Arini et al., 2023); on the other hand microalgae, that are considered to be the most sensitive taxonomic groups and commonly used as indicators of environmental pollution since they are the main primary producers at the basis of the food chain. Any disturbance of the microalgae population will have an effect on the whole ecosystem (Neidhardt & Wasmuth, 2012). More precisely, the organisms were exposed for one week to five different types of NPs (from 1 to 1000 µg/L) made from bioplastics already marketed or still under developement. The algal growth was monitored for one week, and bivalves were tested through filtration tests and gene expression studies. This study aimed to bring the very first results of testing biosourced NPs on freshwater and marine aquatic species and to provide highly innovative insights in order to help use them in an environmentally safe manner.

## METHODS

### Nanoplastic preparation and characterization

Different bio-sourced plastics were used to synthetize Bio-NPs under laboratory conditions: Polycaryophyllene (PCAR) polymer synthesized in-lab from a terpene present in essential oils (Medeiros et al., 2021); Polyamide 11 (PA11, a bio-sourced polymer produced from castor oil commercialized by the Arkema company under brandname Rilsan™); NIPU non isocyanate polyurethane synthesized in-lab by reaction of cyclic carbonates and diamines, derived from pripol and primane provided by Croda) (Maisonneuve et al., 2015; Bobbink et al., 2019); Polyhydroxyalkanoates (PHA) provided by NaturePlast, Caen, France; Polylactic acid (PLA) Ingeo 4032D provided by NatureWorks, Plymouth MN, USA.

PCAR and NIPU pellets were dissolved in tetrahydrofuran solvent (THF purchased from VWR, CAS 109-99-9) (10 mg PS/mL THF) for 10 min under agitation. In order to nanoprecipitate the polymer chains into nanoparticles, milliQ water (10:1 volume ratio) was added under stirring process, and left under agitation for 2 h at 50°C, according to standard solvent-shift procedure as in Schubert et al (2011). As described previously (Arini et al., 2023), the suspensions were left open under a hood for one week to ensure total evaporation of the THF solvent. Previous analyses performed by GC-MS on similar samples confirmed that this evaporation time was sufficient to no longer detect any residual presence of THF into the solution (Arini et al., 2023).

PA11, PLA and PHA pellets were fragmented and sieved as described in Arini et al (Arini et al., 2023) with slight modifications. Briefly, plastics were first crushed in a mill grinder (IKA- A10) filled with liquid nitrogen. Second, the millimetric powders were crushed in a centrifugal grinder (Retsch ZM 200) cooled by continuously pouring liquid nitrogen on the rotor blades with successive sieves of 2 mm, 1 mm, 500 µm, 200 µm, and finally 80 µm threshold size. The finest powders were resuspended into milliQ water and sonicated for 10 min. The suspensions of PA11 and PHA were filtered with teflon membranes (Versapor, pore size of 0.8 µm) whereas for PLA, glass-fiber (GF) syringe pre-filters (Nalgene^®^, pore size of 1 µm) were used in order to obtain nanoscale solutions. In this particular case, 1 g of the PLA powder was first nanoprecipitated from THF into water, then dried overnight to produce a foamy material (as shown on Fig A1 in Annex), which was cryo-milled in a second step. This procedure helped to reach better dispersed state of PLA nanoparticles.

Polymer concentrations in the aqueous suspensions were determined by thermogravimetry analysis (TGA) with a Q500 apparatus by TA Instruments. In practice 100 µL aliquots were heated up to a plateau at 120°C and the concentration was obtained from the dry extract mass. The hydrodynamic size of the particles was measured using both the Cumulant fitting method and the Pade-Laplace multimodal analysis on two dynamic light scattering (DLS) instruments from Cordouan technologies (Pessac, France) operating at backscattering angles of respectively 165° (Vasco™ Flex) and 170° (Vasco™ Kin). Zetametry data were aquired by a phase analysis light scattering (PALS) instrument NanoZS from Malvern Inc. (Malvern, UK) operating at 173° angle to measure the electrokinetic mobility of the nanoparticles and deduce their zeta-potential, which sign and amplitude characterize their surface charge once the NPs are dispersed in water.

### Algal growth experiment

Two fresh-water and one marine algae were tested for the experiment: *Desmodesmus subspicatus*, *Chlorella vulgaris* and *Tetraselmis suecica*. These are unicellular algae, which makes their concentrations easy to estimate by spectrophotometry or counting and assay in growth experiments. Standard curves were established for the three species to correlate optical density (OD, measured with a Jenway^®^ 6300 Visible Spectrophotometer) to each algal concentration. Algal suspensions were prepared in autoclaved media (Dauta for freshwater algae or F/2 for marine one). The day of start, three algal suspensions (for each species tested) at a concentration of 10^6^ cells/mL were prepared. A volume of 90 mL was poured in each corresponding glass vial, and placed on a platform shaker, under light (12:12, 3 replicates per condition) for 7 days.

The five Bio-NPs were assayed (PCAR, PA11, NIPU, PHA, PLA). Three concentrations of each Bio-NP were tested: 10, 100 and 1000 µg/L. We chose to use rather high concentrations since growth was monitored in the very first hours of exposure (from 24 h to one week). Each nanoplastic stock suspension was sonicated for 10 min before being spiked in algae vials. These concentrations were chosen to be as low as hypothetical concentrations of petro-sourced microplastics into the environment (Goldstein et al., 2013; Desforges et al., 2014) and sufficiently high to supposedly trigger effects after a short-term exposure. Each day, vials were randomly moved from the middle to the side of the shacking platform, to prevent from border- line effects with light.

OD was measured in each vial after 24 h, 48 h, and one week of experiment to monitor algal growth among the different conditions tested. Samples were aliquoted periodically to check the cellular numeration under microscopic observation and confirm the validity of the standard curve used. Data corroborated throughout the experiment.

### Corbicula fluminea collection and acclimation

Asiatic clams, *Corbicula fluminea*, were used as filter-feeding bivalves for the experiment. Around a hundred clams were collected in a small tributary of the Isle River in Montpon- Ménestérol. Clams were maintained in 100L tanks filled with clean water for 2 weeks of acclimation, before the start of experiment. They were fed every other day with the green algae *Desmodesmus subspicatus* (70 000 cells/individual/L).

### Clams exposure and filtration rate

At 24 h before the start of the experiment, 1 liter glass units were filled with dechlorinated tap-water aerated with pumps under light / night cycle (12 h / 12 h) and constant temperature (15°C). The day of the start of the experiment, two clams were added per experimental unit. They were exposed to one of the five Bio-NPs tested (PCAR, PA11, NIPU, PHA, PLA) for 7 days. Three concentrations of each Bio-NP were tested: 1, 10 and 100 µg/L. We chose to use lower concentrations than for algae to prevent from potential mortality of bivalves at high concentrations. Since no concentrations are known in the environment for NPs, we chose low experimental concentrations, in the range of µg/L, to be as close as possible to the supposed environmental conditions. Each Bio-NP solution was sonicated for 10 min before being spiked in experimental units. There were five experimental replicates per condition. Water was renewed every other day, and Bio-NPs were added in experimental units right after water renewal to ensure having concentrations as close as possible to nominal ones. During the experiment, clams were fed with the algae *D. subspicatus* (70 000 cells/individual/L) two-hours before water renewal to avoid Bio-NPs’ adsorption onto algae cells and ensure a waterborne exposure. No mortality was observed in clams throughout the experiment.

The seventh day, one clam per experimental condition was retrieved from its experimental unit to perform the filtration test. Each clam was placed in an individual NPs-free vial filled with 50 mL of a *D. subspicatus* suspension with an initial concentration identical in each vial at the start of the filtration test (1.8 10^6^ cells/mL). This concentration was chosen according to previous experiments which concluded that it was suitable to ensure a good production of faeces and pseudofaeces resulting from an efficient feeding of bivalves (data not shown). There were five experimental replicates per condition. OD was measured after 60 and 120 minutes in order to calculate filtration rates of clams, depending on their exposure conditions. The left-over clams not used for the filtration test (one clam per experimental unit) were also placed in 50 mL vials filled with 1.8 10^6^ cells/mL of *D. subspicatus*) to ensure having the exact same conditions among clams throughout the filtration test and before dissections.

Filtration rates (FR, cells/min) were calculated as follows:

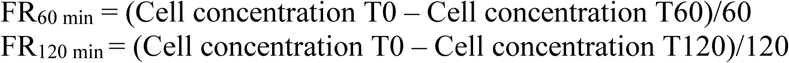

Then, clams were sacrificed to dissect gills and visceral mass for qPCR analyses. Organs from two individuals originating from the same experimental unit were pooled to get enough mass. Samples were stored in RNA later at -80°C before analysis.

### Transcriptomic analyses

Gills and visceral mass were used to perform real-time PCR in order to assess differential gene expression involved in a variety of cellular functions. Total mRNA was extracted using a specific kit (SV Total, Promega) according to the manufacturer’s instructions. The purity (ratio 260/280 >2) and the concentration of total mRNA were checked by spectrometry (BioTek EPOCH with a Take3 plate). The RNA samples were diluted to a concentration of 100 ng/µL. Diluted samples were used for the reverse transcription (RT), to synthetize copy DNAs using a RT system kit (GoScript, Promega) in an Eppendorf Mastercycler™. Samples were kept at -20°C until their use for Quantitative polymerase chain reaction (qPCR) amplifications.

The GoTaq^®^ qPCR Master Mix kit (Promega) was used to perform qPCR amplifications in 384-well plates. Each well was filled with 1 µL of a mix of forward and reverse primer of each gene (5 µM, Table 1), 12.5 µL of the qPCR Master Mix, 6.5 µL of RNA free water, and 5 µL of cDNA. The qPCR program was performed in a LightCycler 480 (Roche) as follows: 95°C for 10 min then 40 cycles at 95°C for 30 s, 60°C for 30 s, and 72°C for 30 s. A dissociation curve was done at the end of amplifications in order to check their specificity. Gene expressions were normalized according to the GeNorm method with two reference genes: *β-actin* and *rpl7* and their differential expressions were calculated with the 2^-ΔΔCt^ method (Livak and Schmittgen(2001)). Induction factors compared to controls were calculated according to the following formula:

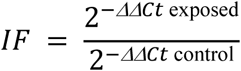

**Table 1:**
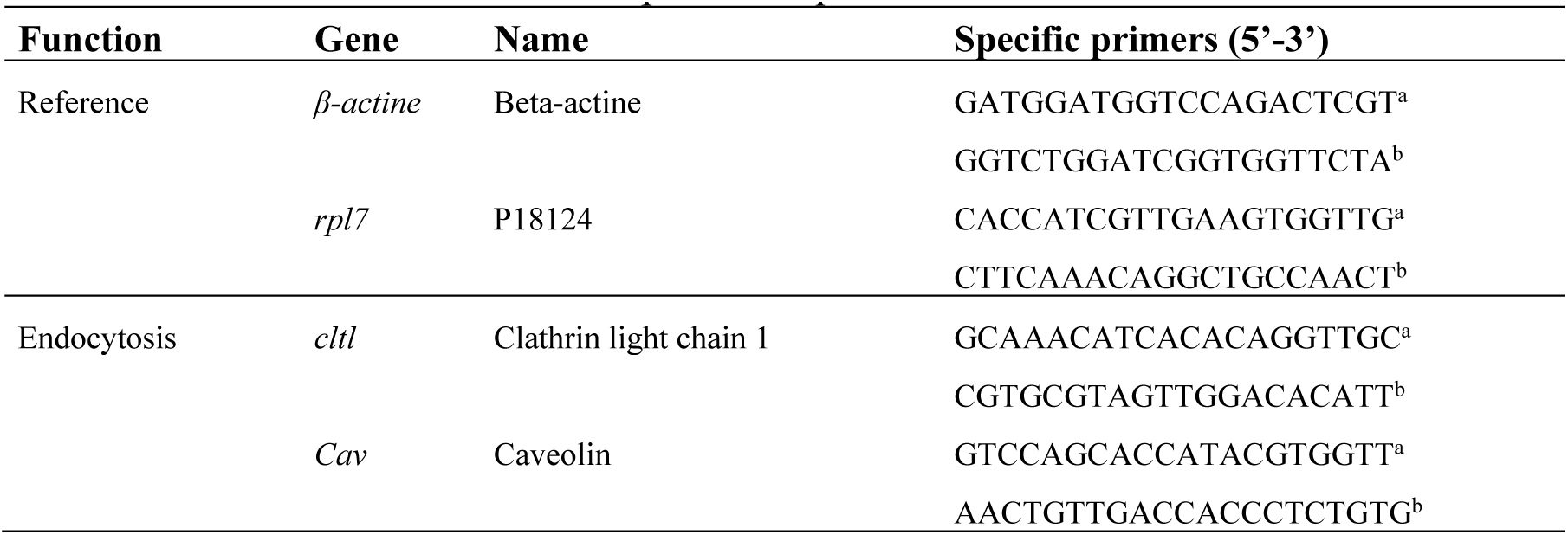
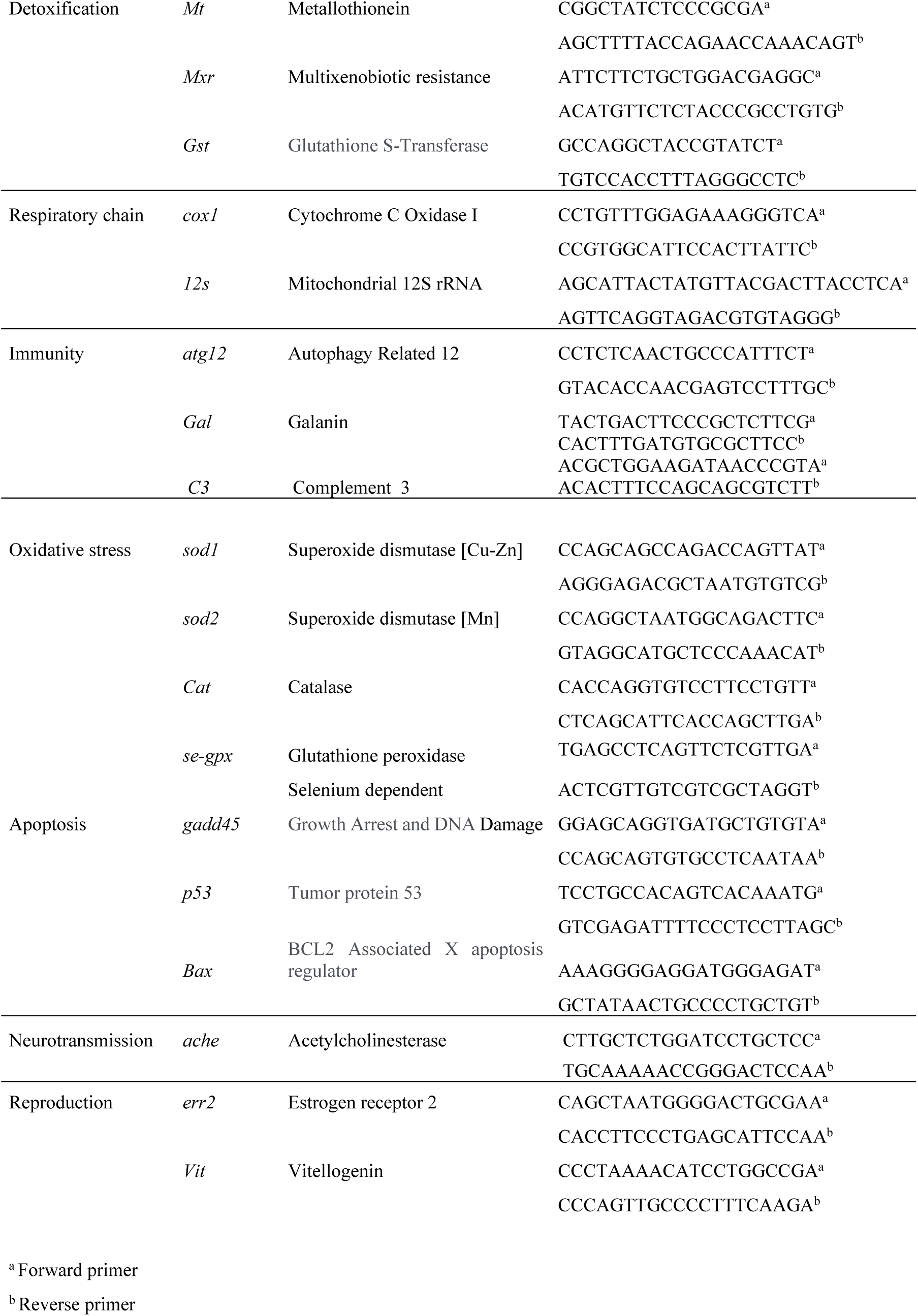
Gene functions studied and primer sequences used.

Twenty genes involved in endocytosis, detoxication, mitochondrial metabolism, immunity, neurotransmission, oxidative stress, apoptosis, and reproduction were assessed (Table 1).

### Data treatment

Non-parametric Kruskal-Wallis tests were used to check significant differences between treatments and controls (XL-Stat software version 2013.5.09, 1995e2013 Addinsoft). p < 0.05 was considered significant for all statistical tests.

## RESULTS

### Bio-NPs characterization

The sizes of the five Bio-NPs were measured by Dynamic Light Scaterring (DLS). Two methods were used to fit the correlograms: Cumulant and Pade-Laplace methods (Table 2), and the mean was calculated to get an averaged size of each NP. The five types of Bio-NPs were in same range of size, comprised between 190 and 322 nm. Table 1 also lists the mean values and standard deviations of Zeta potentials, which are all moderately negative (from -14 to -21 mV) as presumably ascribed to the adsorption of partially hydrophobic anions onto the polymer surfaces, like HCO3^-^ (Yan et al., 2018). The distributions of Zeta potential are displayed in Annex (Fig A2).

**Table 2:**
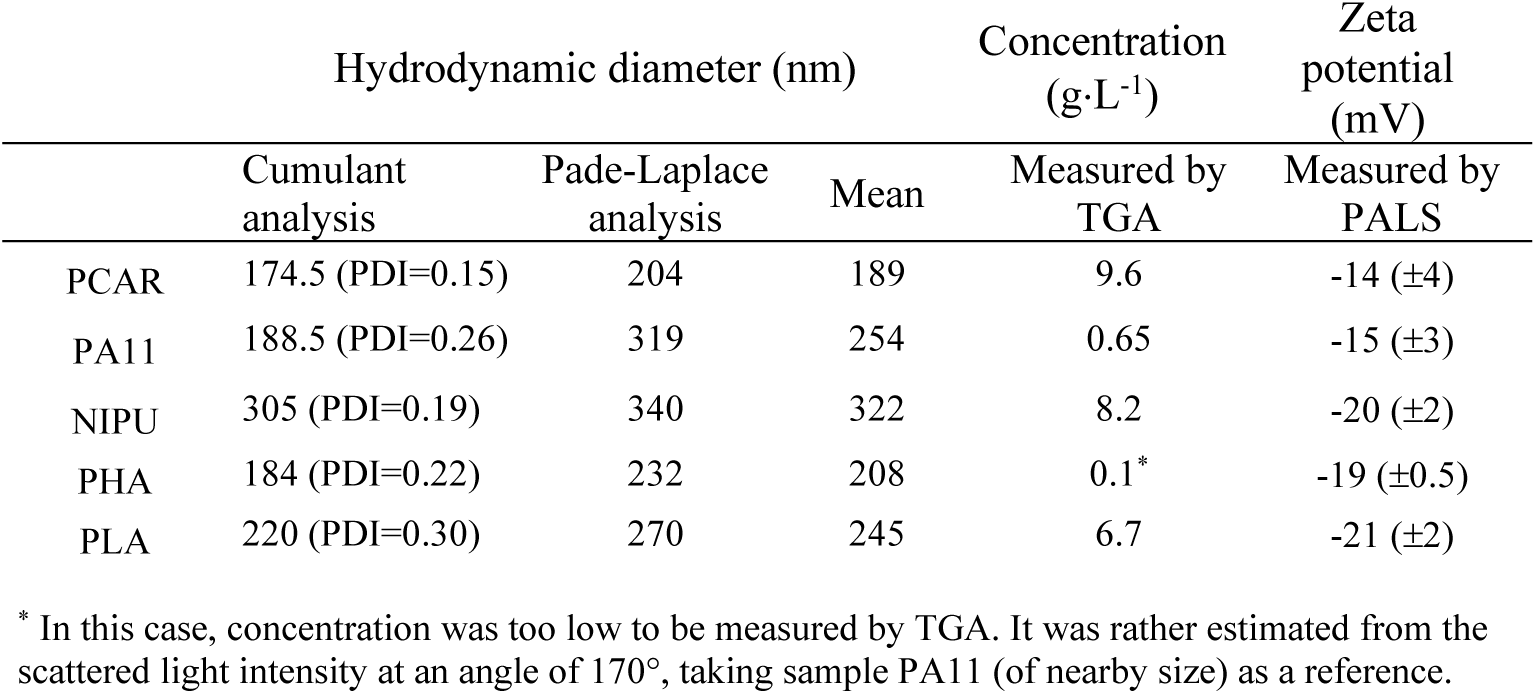
Bio-NP hydrodynamic sizes according to Cumulant (with polydispersity index, PDI) and Pade-Laplace methods, and their mean, values of weight concentration measured by TGA, and zeta potential by PALS in pure water (see distributions in Fig. S2).

### Growth of algae exposed to Bio-NPs

NPs impaired algae growth for each polymer tested, from 1 day to 1 week exposure (Figure 1). PCAR exposures resulted to cell concentration decrease of *D. subspicatus’* at all concentrations tested after 24h and 48h, but algae concentrations were not different from controls after one- week exposure. No effects were observed for the two other algae.

**Figure 1:**
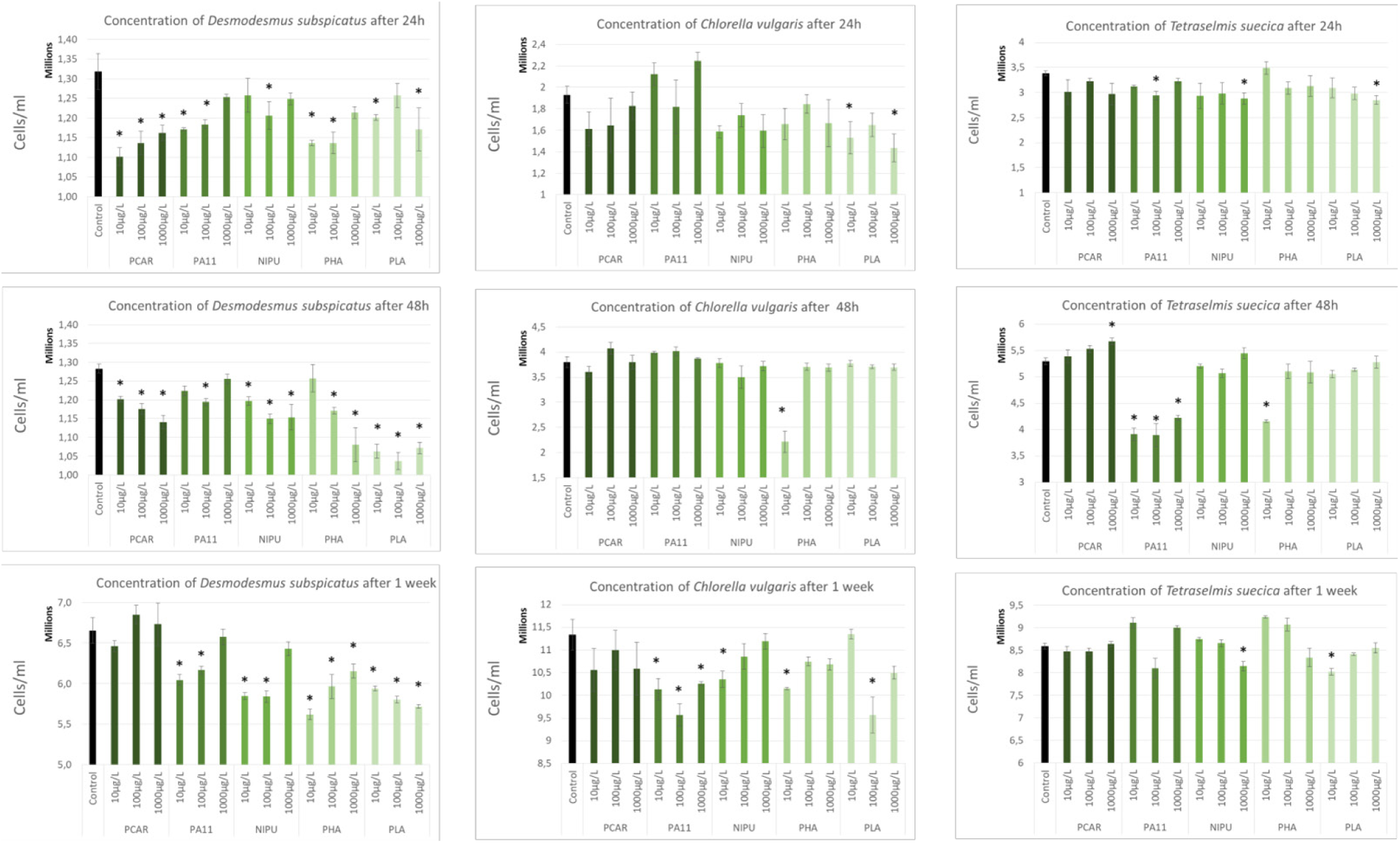
Cellular concentrations (cells/mL) of the three different algal species after 24h, 48h and one week of exposure to the different polymers tested (PCAR: polycaryophyllene, PA11: polyamide-11 (Rilsan™), NIPU: non-isocyanate polyurethane, PHA: polyhydroxyalkanoate, PLA: polylactic acid) at 10, 100 and 1000 µg/L (n=3, Kruskal-Wallis test p<0.05).

PA11 significantly decreased cellular concentrations compared to control from 24h to 1-week exposure in *D. subspicatus* and *Tetraselmis suecica* cultures. Cellular growth was also inhibited in *Chlorella vulgaris* cultures after one week of exposure. It is worth noticing that the highest PA11 concentration (100 µg/L) did not trigger effects on *D. subspicatus* growth.

NIPU exposure provoked few growth inhibitions on *C. vulgaris* and *T. suecica* whereas *D. subspicatus* was sensitive to this polymer. Growth was significantly lower than controls in algae exposed to 10 µg/L of NIPU after 24h, 48h and one week of treatment. The lower condition (10 µg/L) also resulted in growth inhibitions after 48h and one-week of exposure

PHA significantly decreased algal concentrations compared to control in *D. subspicatus* cultures for all times and concentrations tested. Few effects were reported for other algae. Only the 10 µg/L concentration resulted in lower algal concentrations after 48h and one week of exposure for those algae.

PLA also impaired *D. subspicatus’* growth for all concentrations tested from 24h to one week. The exposure to *C. vulgaris* and *T. suecica* resulted in few growth inhibitions after 24h and one week.

Radar diagrams presenting the inhibition rates (%) of algal growth for each polymer and each alga tested (Figure 2) show that the highest growth inhibition occurred in *D. subspicatus* algal units at the lowest concentrations tested (10 µg/L). There was an average of *D. subspicatus’* growth inhibition of 10.1 ± 2.7 % among all NPs tested. Growth inhibition was also important in *C. vulgaris*, and stronger in conditions with 100 µg/L of NPs (mean of 8.7 ± 3.6 %). The lowest effects were observed for *T. suecica* with a growth inhibition lower than 1.8 ± 1.3 % in all conditions tested. The growth inhibition was the lowest with the highest concentration tested (1000 µg/L) for all three algal species.

**Figure 2:**
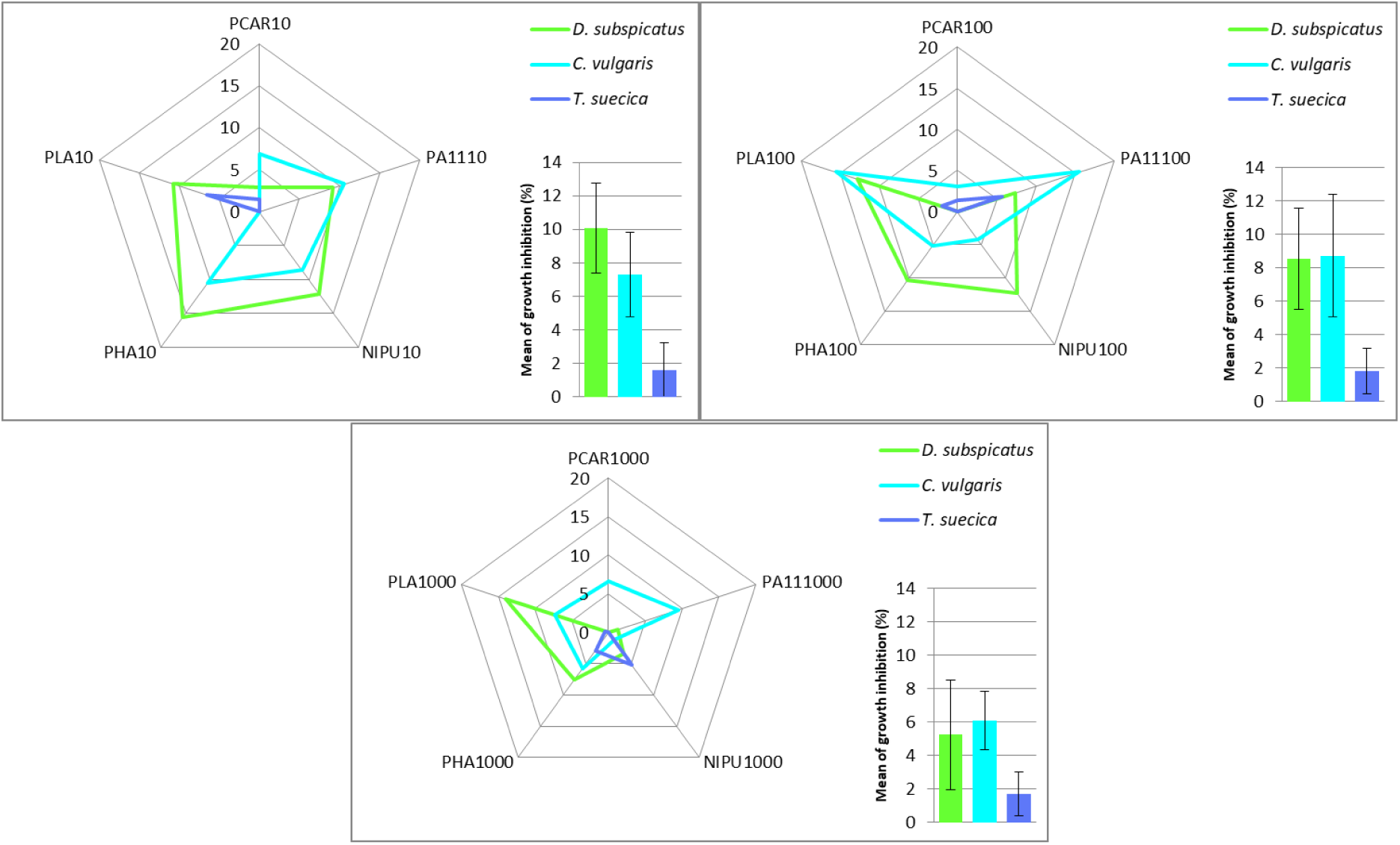
Radar diagram presenting the inhibition rates (%) of algal growth for each polymer and each alga tested at 10, 100 and 1000 µg/L after one week of exposure; histograms: mean of inhibition rates (%, n=5) of all Bio-NPs tested in each species.

PLA is the most deleterious for algal growth with a mean of inhibition among the three concentrations tested of 7.7 ± 1.3 %. PCAR is the less harmful in terms of algal growth with a growth inhibition averaging 2.8 ± 0.3 %. PA11, NIPU and PHA are in between with a mean of inhibition of 6.6 ± 1.7, 5.2 ± 1.1, and 6.4 ± 1.1%, respectively.

### Filtration tests

Each NP tested resulted in significant decrease of the filtration rates of clams after one week of exposure (Figure 3). After 60 min, PHA generated effects with the lowest concentrations tested (1 and 10 µg/L) whereas PCAR, PA11 and NIPU had significant effects with higher concentrations (10 and 100 µg/L). The exposure to PLA resulted in a decrease of the filtration rate for the 10 µg/L condition, only.

**Figure 3:**
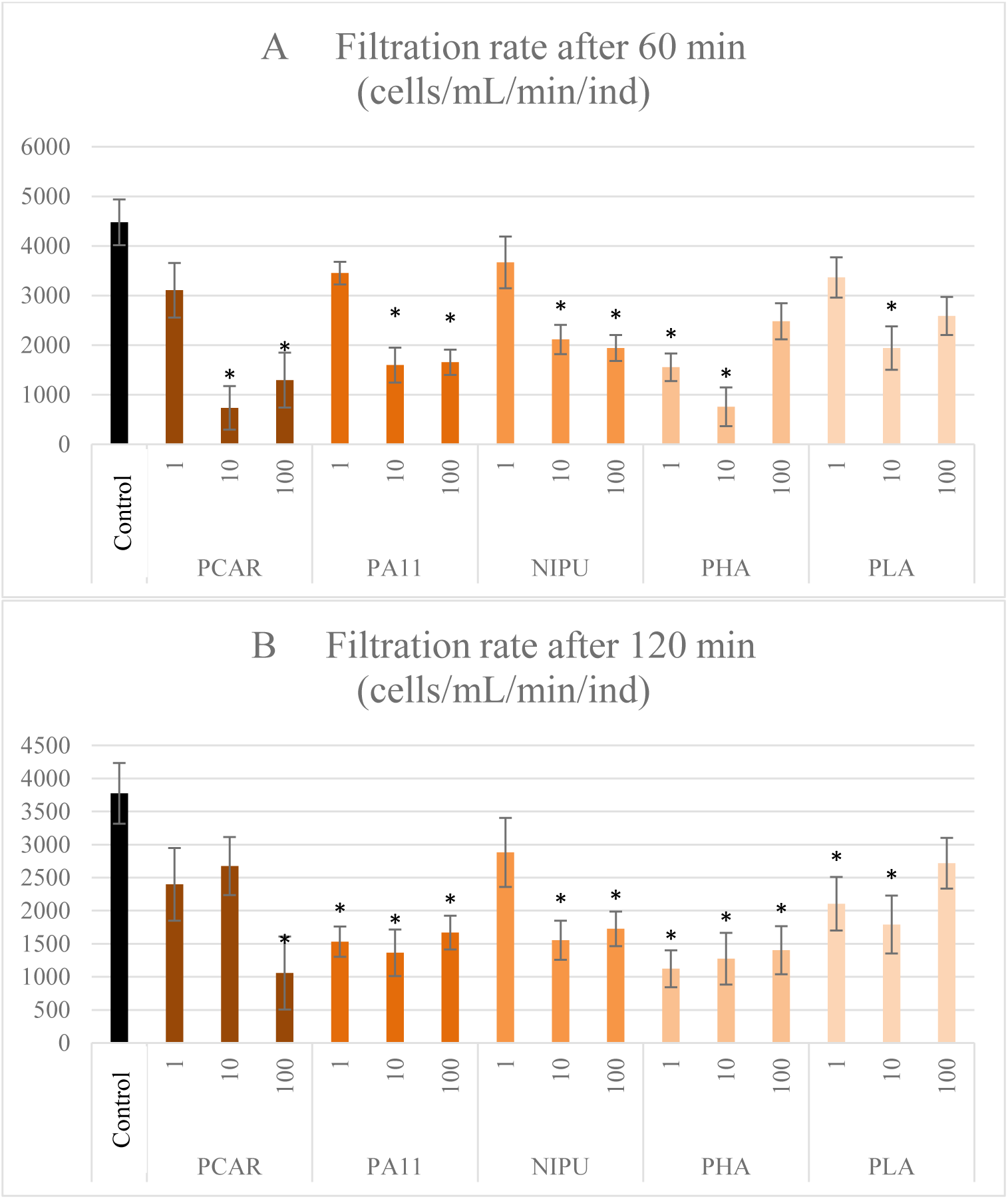
Filtration rates (cells/mL/min/ind) after A: 60 min, and B: 120 min of control *Corbicula fluminea* and of individuals exposed for one week to the different polymers tested (PCAR: polycaryophyllene, PA11: Rilsan™, NIPU: non-isocyanate polyurethane, PHA: polyhydroxyalkanoate, PLA: polylactic acid) at 1, 10 and 100 µg/L (n=5, Kruskal-Wallis test p<0.05).

The effects on filtration were still significant after 120 minutes in NP-free environment. Some effects on filtration rates appeared later than 60 min and were significantly different from control only after 120 min. While PCAR, PA11 and NIPU had only generated effects on filtration rates at 10 and 100 µg/L after 60 minutes, the results after 120 minutes showed that clams exposed to the lowest concentrations (1 µg/L) of PA11 and PLA have filtration rates significantly lower than controls. On the opposite trend, the filtration rates of clams exposed to PCAR at 10 µg/L were no longer significantly different from controls after 120 minutes.

### Relative gene expression Visceral mass

Transcriptomic results obtained in visceral mass of clams showed that each NP tested induced gene modulations in genes involved in all the functions tested (Table 3). Endocytosis and detoxication genes were up-regulated with the five polymers tested. The strongest modulations were observed for PCAR-1 and NIPU-1 (359.9 and 86.4 fold-changes for *clhc*, respectively; and 557.6 and 55 fold-changes for *mxr*, respectively). The mitochondrial metabolism was also affected and the *12s* and *cox1* genes were induced in the PCAR-1, PA11-100 and NIPU-1 conditions, whereas they were repressed in the PCAR-100, PLA-1 and PLA-100 conditions. Immune responses were implemented with the up-regulation of either *atg12*, *gal* or *c3* in all conditions tested, except PCAR-100 and PLA-100 where genes were repressed. Immune response was mainly promoted in PCAR-1 and NIPU-1 conditions (for example 307.3 and 145.3 fold-changes for *c3*, respectively). The acetylcholinesterase gene was strongly induced in the PCAR-1 and NIPU-1 conditions (68.3 and 25.3 fold-change, respectively), and to a lesser extent in PHA-100 condition (4.6 fold-change), and was repressed in PCAR-100 (0.1 fold-change). Each exposure to one polymer led to strong up-regulations of genes involved against the oxidative stress (except for PCAR-100). Among them *cat* and *gst* presented the highest gene expression (1184.7 and 1418.1 fold-change for PCAR-1, for instance). Genes promoting apoptosis were also strongly induced in all conditions tested (except PCAR-100 and PLA-100) especially at low concentrations of NPs (for instance we obtained fold-change factors of 319.9 and 116.9 for *gadd45* for PCAR-1 and NIPU-1, respectively). Genes involved in the reproductions were also impacted with strong up-regulations of *err2* and *vit* (except for PCAR- 100 and PLA-100).

**Table 3:**
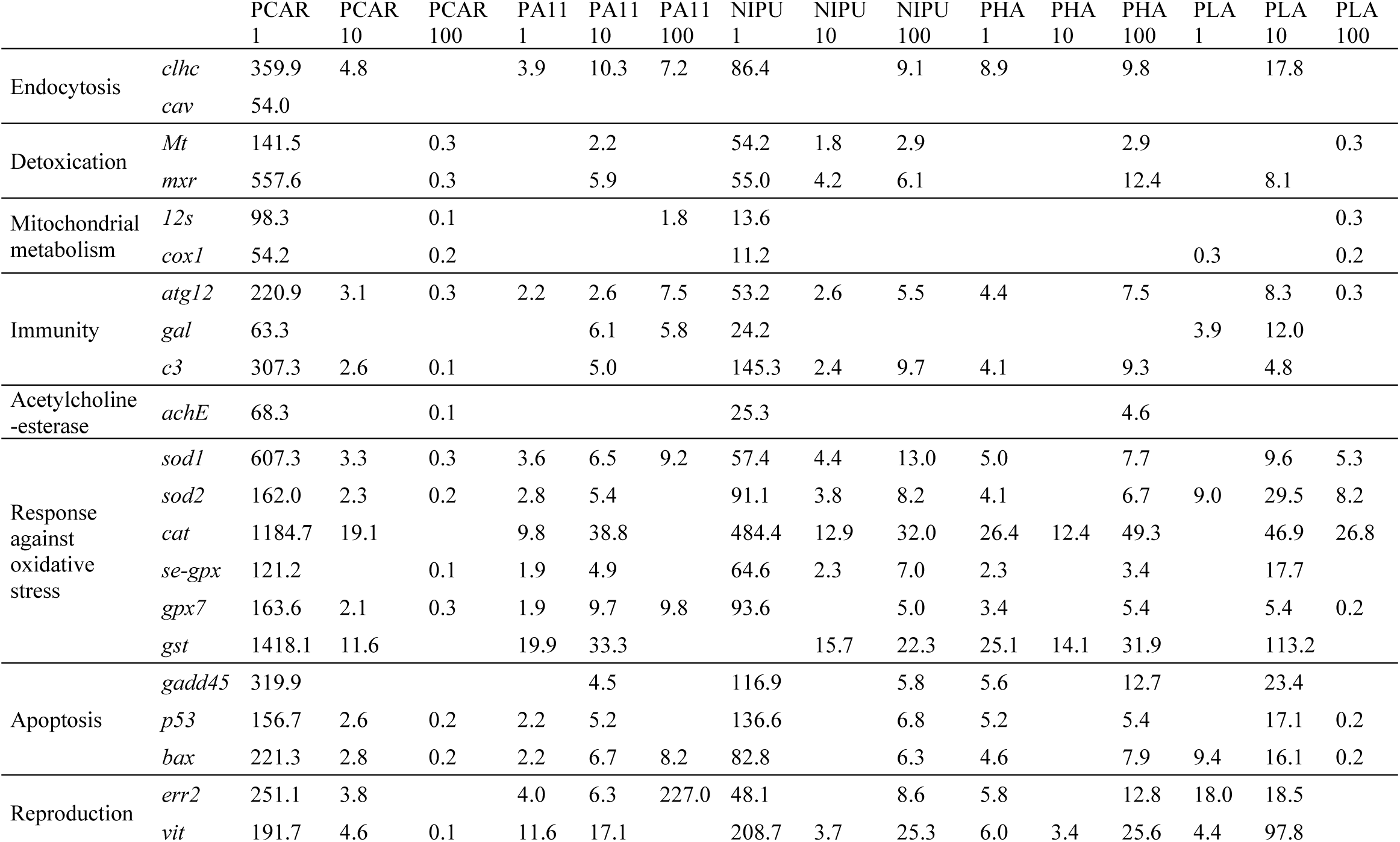
Differential gene expression (mean, n=5) observed in visceral mass of *Corbicula fluminea* exposed to Bio-NPs. Results are given as induction (>1) or repression (<1) factors as compared to controls. Only statistically significant results are reported (Kruskal-Wallis test p<0.05).

Genes were grouped by functions. Fold-change factors were averaged to get a mean value of gene expression among each functional group (as gene expressions were following the same pattern of regulation within each group, either up- or down-regulated). The geometrical mean was chosen in order to get a better estimate of the central value of the data. Results were plotted as radar diagrams, presented in Figure 4.

**Figure 4:**
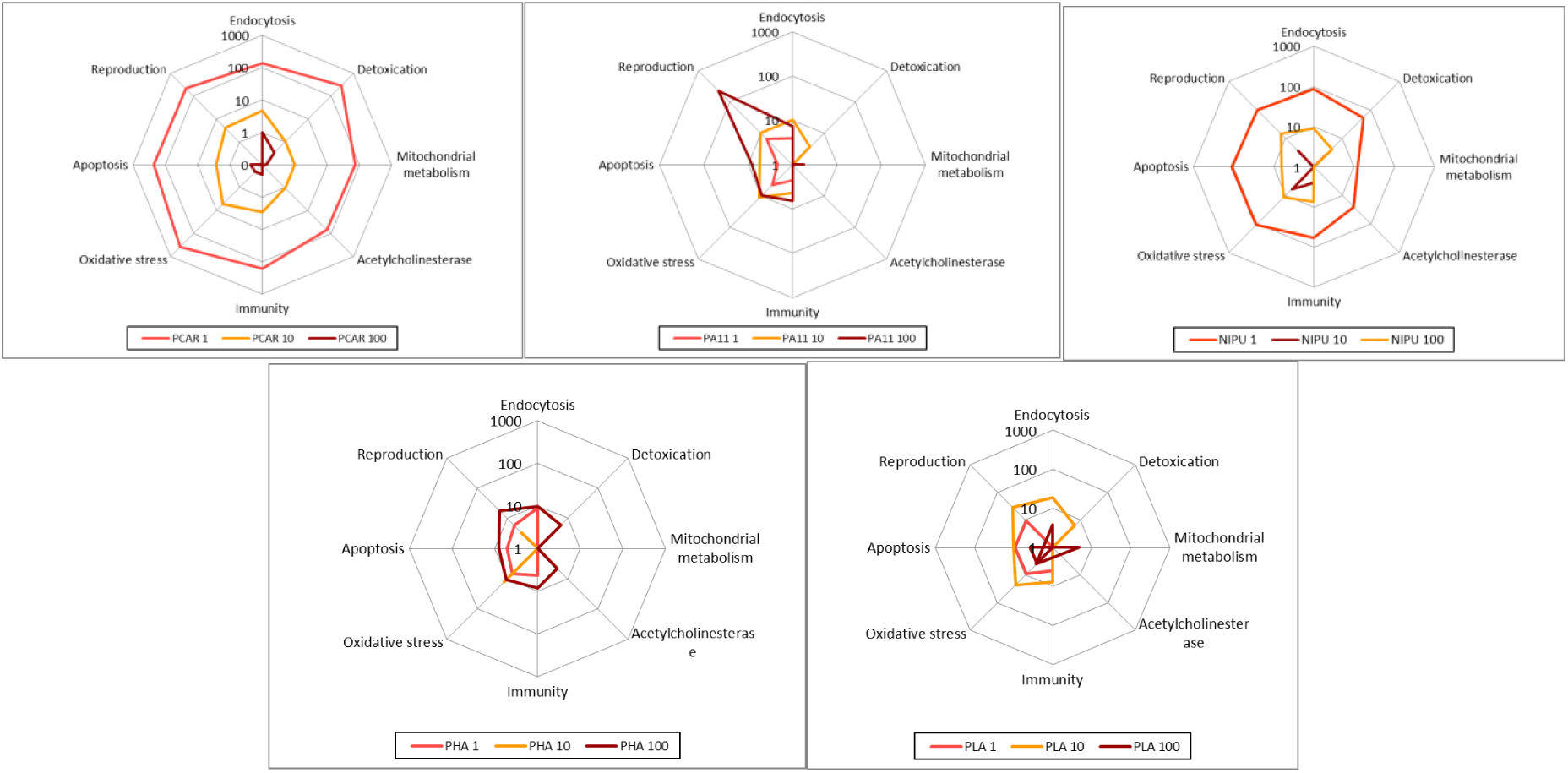
Radar diagrams presenting the average (n=2 to 6 depending on gene function) of fold-change factors among cellular functional groups of visceral mass of *Corbicula fluminea* exposed to the Bio-NPs (PCAR: polycaryophyllene, PA11: Rilsan™, NIPU: non-isocyanate polyurethane, PHA: polyhydroxyalkanoate, PLA: polylactic acid) at 10, 100 and 1000 µg/L.

From those diagrams, results easily show that PCAR and NIPU are the polymers which ended up in the strongest gene modulations compared to other polymers tested. Their lowest concentrations (1µg/L) resulted in stronger effects on gene modulation than higher ones (10 and 100 µg/L). All gene functions tested were impacted by those two polymers at 1 and 10 µg/L but no more at 100 µg/L where effects on gene modulation were strongly reduced.

PLA and PA11 effects are ratherly more pronounced for the 10 µg/L condition, compared to 1 and 100 µg/L. The effects on gene modulations are stronger than PCAR at high concentrations (100 µg/L) and all gene functions tested were impacted among the three NP concentrations tested (except ache for PA1110).

PHA showed lower effects than the other polymers, especially at 10 µg/L. Unlike the other polymers tested, the induction factors were stronger with the highest concentrations tested and effects were not reported for all gene functions tested. Indeed, no gene modulation was observed for the mitochondrial metabolism in PHA conditions.

### Gills

Fewer and weaker gene modulations were observed in gills, compared to visceral mass (Table 4).

**Table 4:**
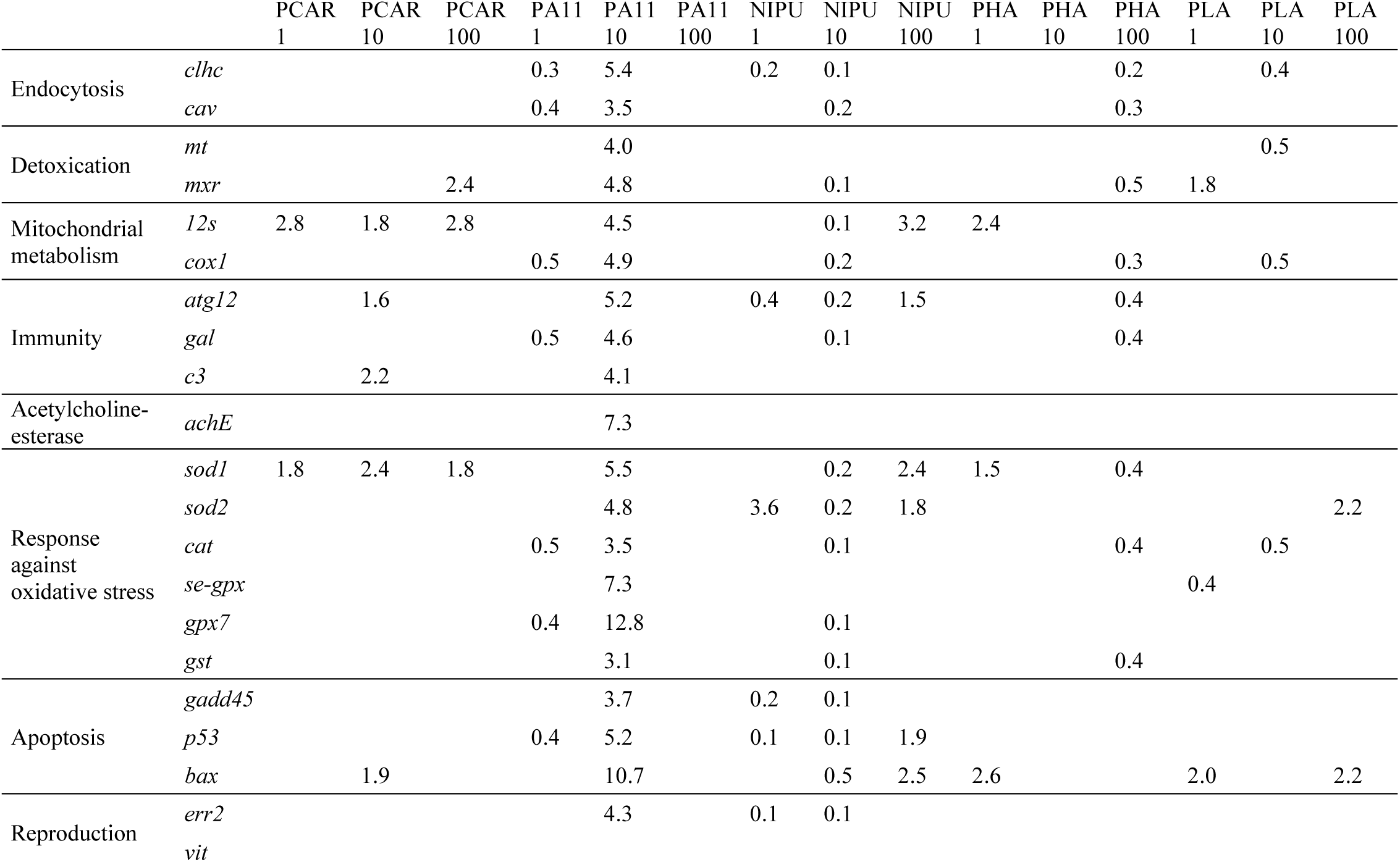
Differential gene expression (mean, n=5) observed in gills of *Corbicula fluminea* exposed to Bio-NPs. Only statistically significant results are reported (Kruskal-Wallis test p<0.05). Results are given as induction (>1) or repression (<1) factors as compared to controls.

PCAR exposure led to an up-regulation of genes involved in detoxication, mitochondrial metabolism, immunity, oxidative stress and apoptosis, but fold-changes were comprised between 1.6 and 2.8. Genes of clams exposed to PA11 (1 µg/L) and NIPU (1 and 10 µg/L) were repressed whereas they were up-regulated for higher concentrations (PA11 10 and NIPU 100). Both polymers resulted in modulations of genes involved in mitochondrial metabolism, immunity, oxidative stress and apoptosis. An opposite trend was observed for PHA whom genes were up-regulated in clams exposed to the lowest concentrations (1µg/L). No significant effects were observed for PHA10 condition, and gene were repressed in the PHA100 condition. All gene functions were impacted except the acetylcholinesterase and the genes involved in reproduction. The effects of PLA depend on the concentration tested, with genes being either up- or down-regulated within the same functional group. We observed for instance the induction of *sod2* (1.2) and the repression of *cat* (0.5) and *se-gpx* (0.4), depending on PLA concentration. No effects on immunity, acetylcholinesterase and reproduction were reported.

Radar diagrams show that, conversely to visceral mass, effects are not dose-dependent, and that gene up-regulations are rather observed with higher concentrations tested. PCAR is the only polymer inducing genes up-regulations in all NP concentrations tested (Figure 5). PLA is the polymer resulting in the most effects at 10 µg/L. Diagrams also clearly show that some gene functions were not impaired by NP exposure, namely endocytosis (PCAR), immunity (PLA), acetylcholinesterase (PCAR, NIPU, PHA, PLA) and reproduction (PCAR, PHA, PLA).

**Figure 5:**
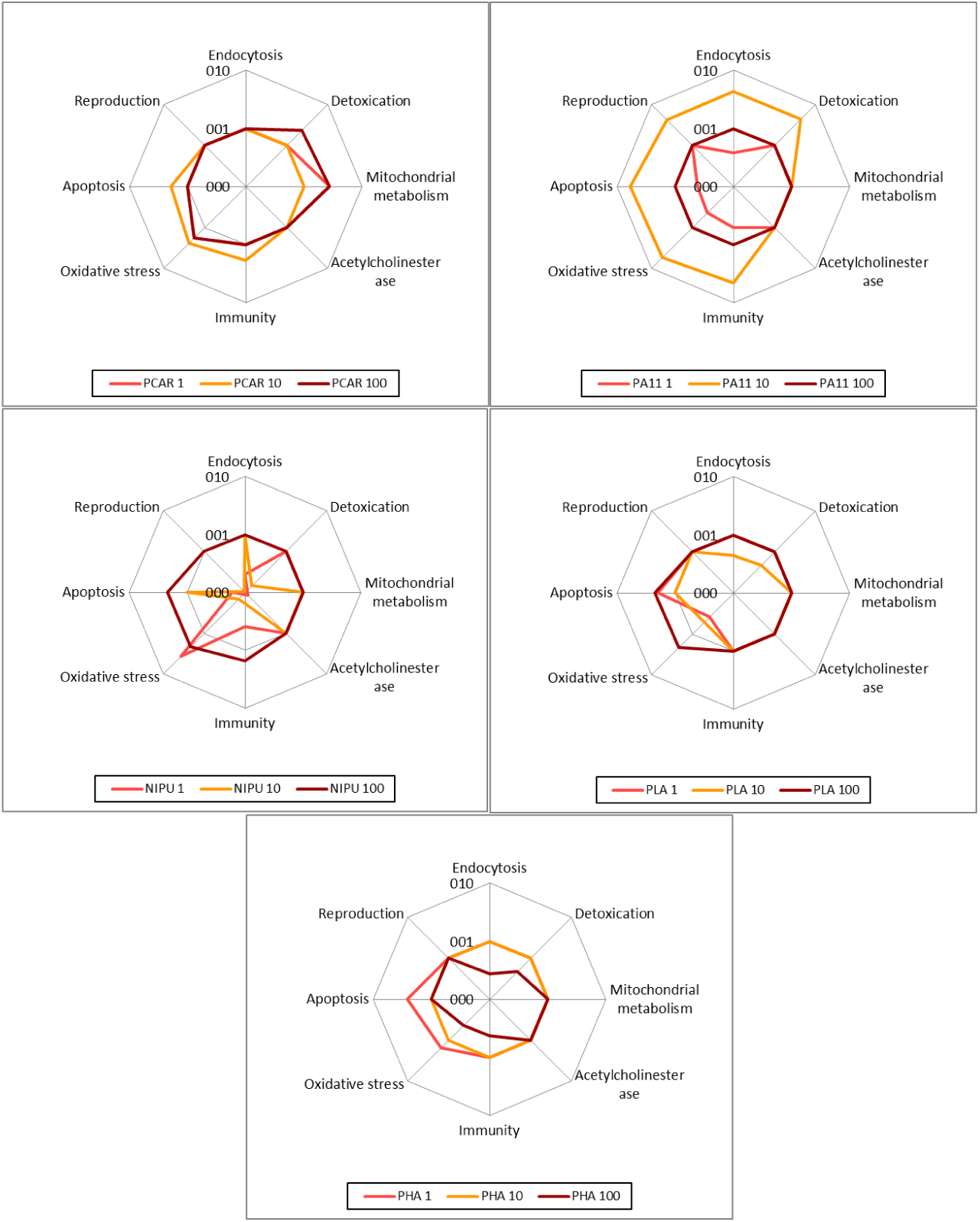
Radar diagrams presenting the average (n=2 to 6 depending on gene function) of fold-change factors among cellular functional groups of gills of *Corbicula fluminea* exposed to the Bio-NPs (PCAR: polycaryophyllene, PA11: Rilsan™, NIPU: non-isocyanate polyurethane, PHA: polyhydroxyalkanoate, PLA: polylactic acid) at 10, 100 and 1000 µg/L.

A scatter plot was made to plot filtration rate and gene expression of *ache* among the different NPs tested (1, 10 and 100 µg/L) for the bivalve *C. fluminea* (after one week of exposure) (Figure 6). A third variable was added on the graph (green circles) which presents the cellular concentrations of *D. subspicatus* depending on exposure conditions (10 and 100 µg/L). The graph shows that all conditions tested ended up in an inhibition of filtration rate, compared to controls. It is worth noticing that the lowest concentrations tested with clams (1µg/L) led to the lowest inhibition of filtration rates (NIPU1, PA11 1, PLA1, PCAR1). PA11 10 is the only condition which induced simultaneously a significant induction of the *ache* gene, filtration decrease, and inhibition of algal growth. NIPU (10 µg/L), PHA (100 µg/L), PLA (10 µg/L) and PA11 (100 µg/L) led to an inhibition of both filtration activity and algal growth, but no significant up-regulation of *ache*. PCAR (10, 100 µg/L) is the only NP tested having no effect on algal growth, and resulting only on an inhibition of filtration rates.

**Figure 6:**
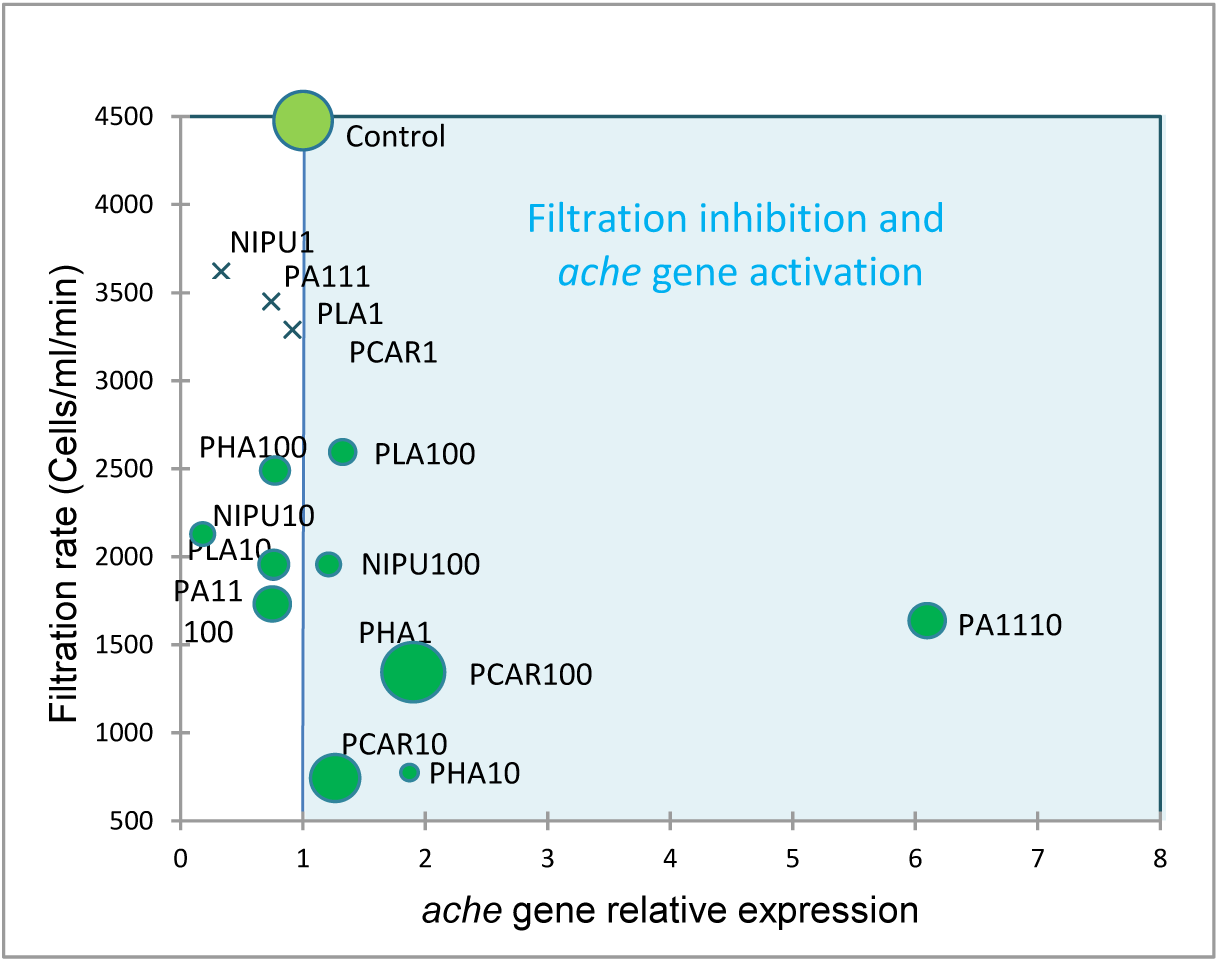
3-factors scatter plot presenting the filtration inhibition rates (cells/ml/min) according to the *ache* gene expression (green circles) of *Corbicula fluminea* exposed to the different conditions for one week. The sizes of green circles represent a third dimension which corresponds to algal concentrations of *D. subspicatus* at 10 and 100 µg/L of NPs (the highest the concentration is, the biggest the circles are). Green crosses replace green circles for concentrations of Bio-NPs not tested with algae (1 µg/L).

## DISCUSSION

Most of the studies report data on petroleum-based nanoplastics or on bio-based microplastics. To the authors’ knowledge, this work is the very first study reporting ecotoxicological data on four species, among animal and vegetal species, from freshwater and marine environments for 5 bio-sourced nanoplastics. We chose algae positioned at the bottom of food-chain, and bivalves, which are filter-feeding animals and act as sentinel species due to their high capacities to accumulate great amounts of xenobiotics from their surrounding environment.

The results presented here deal with critical endpoints for species health and survival, at the individual, cellular and molecular levels (behavior, growth, gene expression). Molecular analyses on gene expressions targeted crucial cellular functions involved in endocytosis, detoxication, mitochondrial metabolism, neurotransmission, oxidative stress, apoptosis and even reproduction. To date, all publications based on bioplastics use high concentrations, in the range of mg/L (González-Pleiter et al., 2019; de Oliveira et al., 2021; Malafaia et al., 2021; Gálvez-Blanca et al., 2026), which is relevant for MPs (Nolte et al., 2017) but probably not for Bio-NPs, or at very short term of exposure (Mustapha et al., 2025). So far, no data exist about concentrations of NPs in the natural environment, and even less concerning concentrations of Bio-NPs. However, the concentrations used in the literature (mg/L) are likely to be much higher than the real concentrations encountered in the environment. In this study we chose to use a gradient of low concentrations, in the range of µg/L (from 1 to 1000 µg/L) to get an overview of Bio-NPs potency to generate toxicity on aquatic organisms, even at very low concentrations (1 µg/L), and after 7 days of exposure.

As Bio-NPs used in our study originate from crushed pellets of raw materials, they differ from final bioplastic products resulting from complex compounding of the polymer with plasticizers or antioxidants. Once the chemical structure of the polymers is fixed, their potential tocicity is also different from the one of the natural products they are originating from, for example fermented cane sugar for PLA, ricinoleic acid for PA11, clove extract for PCAR, fatty acid based polycarbonates and amines for NIPU… Assuming that the virgin polymers used in this study contain only low levels of additives, the potential toxicity arised from the Bio-NPs themselves.

### Bio-sourced nanoplastics generated an inhibition on algal growth

Bio-NPs were inoculated in unicellular algal cultures to assess their impact on cell growth. We chose nanoplastics made from different natural bio-sourced polymer materials, with similar sizes and zeta potentials to have homogeneous characteristics among them.

All five Bio-NPs tested generated growth inhibitions in at least one of the tree algae tested. The freshwater species *D. subspicatus* was the most sensitive to Bio-NPs, and showed the earliest effects compared to the other algae, especially at low concentrations (10 and 100 µg/L). The five Bio-NPs tested led to growth inhibitions of this species. *C. vulgaris’* growth was also inhibited but effects appeared mainly after one week of exposure for all Bio-NPs tested (except PCAR).

PLA and PA11 were the most deleterious for algal growth among the five Bio-NPs tested, whereas PCAR was the less harmful. It led to growth inhibitions only in *D. subspicatus’* cultures after 24 and 48 h, and no more effects were observed after one week of exposure. It suggests that PCAR interacts in the short term on algae and to a lesser extent in the longer term. Others Bio-NPs tested generated early effects in *D. subspicatus* (after 24h) which were still present after one week of exposure.

Only one study testing Bio-NPs on algal growth has been published so far in the literature. González-Pleiter et al (2019) have reported growth inhibitions on the freshwater algae *Chlamydomonas reinhardtii* after a 3-day exposure to PHB-NPs (PHAs’ family) at 50 mg/mL, concomitant with an increase of intracellular levels of reactive oxygen species (ROS). However it is worth noticing that authors used very high concentrations compared to our experimental conditions. In our study we observed that growth inhibition was lower at high concentrations (1000 µg/L) compared to lower concentrations (10 and 100 µg/L). It has been shown that NPs aggregate at high concentrations (Gigault et al., 2018) and become less bioavailable to aquatic organisms (Hotze et al., 2010), especially in salt water. It suggests that effects may be underestimated when using high concentrations (in the range of the mg/L) or marine waters.

Other studies have been conducted on bio-sourced plastics but dealing with micrometric scales and not nanoplastics, and still at high concentrations (in the range of mg/L). For instance, a study showed that a 11-day exposure to microplastics (MP) of PLA (100 mg/L, 57 µm) inhibited *Chorella vulgaris’* growth and they suggested that effects could be attributed to physiochemical properties of MPs such as the release of additives or shading effects (Su et al., 2022), which would in turns lead to a decrease of photosynthesis and then a reduced fitness of algal population and growth. Indeed, a study on polystyrene nanoplastics (PS-NPs, 1.8–6.5 mg/L) on *D. subspicatus* and *C. vulgaris* cultures resulted in an inhibition of photosynthesis, which was supposedly due to shading effect of the adsorbed PS beads and/or due to obstructed CO_2_ gas flow and nutrient uptake pathways (Bhattacharya et al., 2010).

Other algal growth inhibitions have been documented for petroleum-based NPs. For instance, Baudrimont et al (2020) showed that cultures of the freshwater algae *Desmodesmus subspicatus* and the marine algae *Thalassiosira wessiflogii* presented lower cell concentrations than controls after being exposed for 48 h to NPs made from reference polyethylene or polyethylene debris (0, 1, 10, 100, and 1000 μg/L). The growth inhibition was stronger in the freshwater species *Desmodesmus subspicatus,* like in the present study. The authors suggested that the difference could either be linked to the diatomic frustul of *Thalassiosira wessiflogii*, which would act as a physical barrier against pollutants, or either to the salted medium of the marine species, whom salinity would enhance the NP aggregation (Gigault et al., 2018; Arini et al., 2022) and result in a lower bioavailability of NPs (Hotze et al., 2010).

In our study, very few effects were reported on the marine green algae *Tetraselmis suecica*, compared to the two freshwater species. Then, in our study the salinity of the medium seems to be an important factor influencing the toxicity of bio-NPs. But other factors may interfere with NPs toxicity. Indeed, it has been proven that the cellular membrane of microalgae is composed of carboxyl and phosphate groups, giving them a negative charge, and that the binding of MPs and NPs onto the cell surface of algae is precisely dependent on their surface charge (Kim et al., 2006). Bhattachyra and al (2010) exposed both *Desmodesmus subspicatus* and *Chlorella vulgaris* algae to PS-NPs and assessed how surface charges of NPs could influence their adsorption onto algal cell surface. The authors demonstrated that *D. subspicatus* and *C. vulgaris* display a zeta potential around -11 and -26 mV, respectively, therefore positively-charged NPs (e.g. bearing amine moieties) are more likely adsorbed on the negative surface of algae than negatively-charged NPs. They also showed that *D. subspicatus* was more sensitive than *C. vulgaris* to PS-NPs. This sensitivity was attributed to algal morphologies, as *D. subspicatus* presents a rough shape thus higher total surface/volume ratio (3-78 × 2-10 µm) than *C. vulgaris* (2-10 µm) (Bhattacharya et al., 2010). Moreover, *D. subspicatus* possesses two pairs of flagella for its motility. The higher surface contact and motility could then explain why *D. subspicatus* presented higher growth inhibitions in our study, compared to *C. vulgaris* and even *T. suecica*, which is much bigger in size (10 µm long × 14 µm large) and presents therefore a lower surface/volume ratio than both freshwater algae. Its lower sensitivity could also be linked to its surface charge (zeta potential unknown) that would confer it a lower affinity for the Bio-NPs which all have ζ potentials from -15 to -20 mV (Table 2).

### Bio-sourced nanoplastics induced gene expression modulations

The results from gene expressions in *Corbicula fluminea* showed strong inductions of all gene functions tested in gills and visceral mass. Our results pointed out that Bio-NPs triggered endocytosis mechanisms, attesting their uptake into the cells, as well as detoxification induction to reject them from cells. They strongly disturbed the mitochondrial metabolism, which resulted in a response against oxidative stress and a stimulation of the immune system. Our results also showed an induction of apoptosis which highlights that protective mechanisms implemented against Bio-NPs may not be sufficient and could lead to cell death. Bio-NPs also acted on genes involved in the neurotransmission and reproductive pathways, which could have generated effects not only at the molecular levels but also at the individual (behavior) and population (reproduction) levels.

PA11 10, NIPU10 and PHA 100 triggered the strongest gene regulations in both organs, but it is worth noticing that effects were stronger in the visceral mass than in gills. Genes were mainly up-regulated in visceral mass, whereas they were repressed in gills of clams exposed to NIPU10 and PHA100.

In the visceral mass, PCAR and NIPU showed the most effects. Gene inductions were stronger at 1 and 10 µg/L when compared to the 100 µg/L condition. PHA is the only Bio-NP showing stronger effects at 100 µg/L. Previous studies on *C. fluminea* exposed to oil-based Petro-NPs for 7 to 21 days resulted in few significant gene modulations after one week (Arini et al., 2023). Only two genes (endocytosis, and acetylcholinesterase) were significantly modulated in clams exposed to NPs (1 and 10 µg/L) (Arini et al., 2023). Effects on gene expressions were mainly reported after 21 days of exposure to Petro-NPs. The authors also concluded that gills were less impaired at gene levels than visceral mass. It is likely that Bio-NPs can decompose faster than Petro-NPs, and molecules they are made of could be more bioavailable once ingested by organisms. Therefore they may interact in the shorter term with key molecular functions.

The main differences in terms of gene modulations between the five Bio-NPs tested could rely on their capacities to be biodegraded once ingested. To the one hand, biodegradability depends on several factors (bacteria, temperature, etc. (Atiwesh et al., 2021)) and may take longer for some Bio-NPs compared to others. On the other hand, bioplastics are made from oils, cellulose, or other natural products, which despite their natural origin, reach concentrations much higher than the ones encountered into the natural environment once they enter aquatic organisms and cells (Malafaia et al., 2021). Those two elements could explain the difference in terms of toxicity observed between bio-NPs.

To date, no study in the literature has reported effects of Bio-NPs at the gene level in aquatic organisms. Our data is the first insight showing that Bio-NPs can act on key molecular events after a one-week exposure, such as oxidative stress and at very low doses. Intracellular ROS production has yet been reported by González-Pleiter et al (2019) who exposed *Daphnia magna* to PHB-NPs (50 mg/mL) for 48 h. These authors showed that associated contaminants of PHB (retrieved after ultrafiltration) did not lead to adverse effects on crustaceans and that effects were truly triggered by NPs themselves. This statement corroborates our findings. Indeed, as mentioned above, our study reports effects of Bio-NPs made from virgin polymers, which do not contain additives, and exclude any synergic effects due to chemicals taking part of their composition.

Data about Bio-NPs are extremely scarce, however an oxidative stress has already been reported for MPs made from bio-sourced plastics. For instance, Malafaia et al (2021) studied the effects of PLA-MPs on tadpoles exposed for 14 days at 760 μg/mL and 15 020 μg/L. They showed that PLA-MPs generated an oxidative stress, resulting in an increase of intracellular ROS production, as well as Catalase and GST activities. Another study has also focused on *in vitro* screening tests to assess the effects of 27 bio-sourced plastics (MPs and NPs mixed) on different cellular functions (Zimmermann et al., 2020). The authors highlighted adverse effects at different levels including endocrinology, oxidative stress, and reproduction through estrogen and androgen receptors. Zimmermann et al (2020) showed that PLA could activate a response towards oxidative stress but not PHA. The study also reported effects on andro- and estrogenic receptors for PLA. In our study, Bio-NPs, including PLA, altered gene expressions of vitellogenin and estrogen err2 receptor. Both were induced in visceral mass after exposure to any of the five Bio-NPs tested. However, Zimmermann et al (2020) attested that effects on andro- and estrogenic receptors were mainly due to additives presents in the bioplastics they tested. They concluded that final products made of bio-sourced plastics may contain complex mixtures of chemicals which were in many cases more harmful than bioplastics made from virgin pellets. Our study showed that virgin Bio-NPs can also be harmful on definite endpoints. Their industrial use, adding chemical mixtures, may lead to even more toxic effects since they may degrade faster and be more bioavailable to aquatic organisms than petroleum-based plastics. But the *in vitro* study over-mentioned targets initial events occurring during the early cellular response, while our study is the first one showing the impairment of major physiological mechanisms, such as algal growth or clams filtration rate, when exposed to five different Bio-NPs.

So far, acute toxic mechanisms of Bio-NPs are still unknown and it is not possible to fully understand how they act on gene modulation, to impair physiological functions and behavior of aquatic organisms. Further investigations are needed to deeply study their toxicity pathways and further explored the deciding factors inducing adverse effects.

### Bio-sourced nanoplastics impaired filtration activity of clams

Clams exposed to Bio-NPs have all shown reduced filtration rates compared to controls, and no return to control levels after 2 h in clean water. No such data exists so far on Bio-NPs effects, but behavioral effects have been reported in other species exposed to MPs of bio-sourced plastics. For instance the swimming speed and distance was reduced in fish larvae of *Danio rerio* exposed for 5 days to 3 and 9 mg/L of PLA MPs (de Oliveira et al., 2021). It was showed an increase of the acetylcholinesterase activity in those fish larvae which could explain the decrease of locomotion, because of its key role in neurotransmission and motor activity. Di Giannantonio et al (2022) also reported behavioral alteration in jellyfish exposed to PLA MPs (1, 10, 100 mg/L) for 24 h which presented reduced frequency of pulsation compared to controls. Other behavioral impairments have been observed on terrestrial animals. Liwarska- Bizukojc (2022) has shown that earthworms exposed to PLA MPs migrated downwards to the bottom soil zone, to avoid plastic exposure. However, all these studies deal with MPs and concentrations in the range of mg/L. Our results provide the first evidence of the adverse effects of Bio-NPs at environmentally relevant concentrations.

Behavior has been assessed in bivalves after petroleum-based NPs exposure. Baudrimont et al (2020) conducted filtration tests on *Corbicula fluminea* but no significant effects were reported after 48 h of exposure to Petro-NPs at 1000 µg/L. However, clams produced more pseudo- faeces than controls when exposed to Petro-NPs and authors suggested that it could result from a protective mechanism intending to reject NPs from the clams’ body, before being ingested. It is possible that at lower concentrations (1, 10, 100 µg/L, such as in our study), which possibly corresponds to lower aggregation rates, these mechanisms are not implemented and NPs are actually ingested, provoking adverse effects such as lower filtration activities. Indeed, previous studies on *Corbicula fluminea* exposed to lower concentrations (1 µg/L) of Petro-NPs made from plastic wastes have resulted in an inhibition of filtration activities (Arini et al., 2023). This statement was concomitant with the induction of the *ache* gene in visceral mass of clams after 7 days of exposure. Our results showed an induction of the *ache* gene only in the PA11 10 condition. No effect on gene expression was observed for other conditions, suggesting that other mechanisms could interfere with the filtration activity of clams exposed to Bio-NPs.

### Comparison of effects generated by different kinds of bio-sourced nanoplastics

The Scatter plot sums up the data about *D. subspicatus’* growth, Corbicula’s filtration rate and *ache* gene expression (Figure 6). It shows that all Bio-NPs tested led to a decrease of filtration rates and an inhibition of algal growth (except PCAR). It does not enable to create clusters of Bio-NPs and shows that effects tested in our study are clearly depending on the bio-sourced plastic itself, the concentration and the endpoint tested. It outlines the complexity of interpreting the toxicity of Bio-NPs and extrapolating it to natural conditions. The present study brings brand new findings and will help better understand impacts of such NPs on aquatic species, in order to get new insights for environmental pollution.

The heatmap of Table 5 representing results from algae and clams (mean of the three concentrations tested for each Bio-NP) clearly shows that all five Bio-NPs tested impaired algal growth and clams behavior and gene expressions. PCAR had very low effects on algae but triggered the strongest effects on clams, namely on oxidative stress and apoptosis. PLA was the most harmful Bio-NPs on algal growth and resulted in strong gene modulations on clams. NIPU had stronger effects than PLA on clams but not on algae. PHA presented the lowest effects on both biological models.

**Table 5:** Heat-map of algae results (in shades of green: growth rate, %) and *Corbicula fluminea* results (in shades of red: filtration rate, %) and gene expression of main cellular functions (fold-change) for the five Bio-NPs tested (mean of the three concentrations tested for each Bio-NP after a one-week exposure).

|  | Algal growth rate |  |  | *Corbicula fluminea* |  |  |  |
| --- | --- | --- | --- | --- | --- | --- | --- |
|  | *Desmodesmus* | *Chlorella* | *Tetraselmis* | Filtration rate | Immunity | Oxidative stress | Apoptosis |
| PCAR | 0.91 | 0.95 | 0.99 | 0.5 | 55.2 | 130.3 | 75.3 |
| NIPU | 0.94 | 0.88 | 0.99 | 0.5 | 22.3 | 42.2 | 39.0 |
| PLA | 0.87 | 0.92 | 0.97 | 0.6 | 4.0 | 11.8 | 6.6 |
| PA11 | 1.00 | 0.95 | 1.02 | 0.4 | 4.4 | 8.4 | 5.3 |
| PHA | 0.89 | 0.93 | 1.03 | 0.3 | 4.5 | 10.3 | 4.8 |

The differences in terms of Bio-NPs effects could be due to several factors: 1) their chemical structure, which influences their capacity to bind to cell walls and to have shading effects and/or to be internalized, 2) their degradation potential and thus their bioavailability into the surrounding environment, organisms and cells, 3) their biodegradation potential once ingested.

Our study shows different trends on algae and *Corbicula fluminea*. The most toxic Bio-NPs on algae are not necessarily the most toxic on bivalves, and *vice-versa*. A mechanical effect may be at the origin of differences observed between algae and clams. Indeed, our study on algae suggests that there can be a shading effect as well as a depletion in gas and nutrient exchanges which lead to algal growth inhibitions (Su et al., 2022). Toxic effects reported on *Corbicula fluminea* (filtration and gene expression) suggest that Bio-NPs may be recognized as organic carbonaceous material (Malafaia et al., 2021) and therefore less rejected through pseudo-faeces than petro-NPs. The intrinsic chemical structure and biodegradation potential of each Bio-NP tested could explain the differences observed in terms of toxicity among algae and bivalves.

## CONCLUSION

This study was conducted in a context of a fast growing production of bio-sourced plastics as an alternative to petroleum-based plastic. No other ecotoxicological study has been conducted so far on the potential toxicity of Bio-NPs. It was expected that their effects would be minimized compared to petroleum-based NPs due to their natural composition. However, our study reveals the very first results showing that Bio-NPs are harmful for aquatic organisms and must be considered with attention in ecotoxicological studies. The bio-sourced plastics tested in our study showed deleterious effects on algae and bivalves after a short-term exposure. It is possible that they are not more toxic chemically speaking than the other types of NPs, but they could cause the same mechanical disorders such as shading and obstruction effects on algae. However, our study on clams clearly showed toxic effects on filtration and gene expressions. The natural origin of components of Bio-NPs could make them more easily ingested by bivalves than Petro-NPs. It would be interesting to assess their degradation capacity and bioavailability once internalized, since it can strongly influence their toxicity. Indeed, it is possible that this early response does not last, due to the rapid and effective degradation of Bio-NPs, once they are broken down into the body. Therefore, it is important to conduct further studies to know if the effects of Bio-NPs are still harmful in the long term. Their effects may disappear faster than Petro-NPs which are scarcely degradable. It is also possible that Bio-NPs do not accumulate as much as Petro-NPs in the aquatic environment due to their higher degradation capabilities. As a consequence, the concentrations encountered by aquatic organisms may be lower than the increasing ones of Petro-NPs. This study is very preliminary but it already points out that bio-sourced plastics must be considered with caution in an alternative approach to petroleum-based plastics. Moreover such evaluation should be implemented in the design of new “green” and bio-based polymers and plastics material in a safe by design approach.

## Aknowledgments

We want to thank the LabEx COTE funding for supporting the PLASCOTE project (ANR 10- LABX-0045) and the University of Bordeaux for funding through its “inter-department” call 2021 (Environmental Sciences and Materials and Radiation Sciences research departements) . We also thank the molecular biology platform of the EPOC laboratory for giving us access to the transcriptomic and enzymatic assays and PCR equipment.

## Open Data

All datasets for Figures and Tables are available on https://recherche.data.gouv.fr repository: A. Arini, A. M. Meideros, V. Coma, E. Grau, O. Sandre, M. Baudrimont, 2026

*All data related to the figures and tables in the article "Are bio-sourced nanoplastics inert for aquatic species? A toxicity study on three micro-algae species and a freshwater bivalve"* https://doi.org/10.57745/FKGAPY, Recherche Data Gouv.

## ANNEX

**Fig A1:**
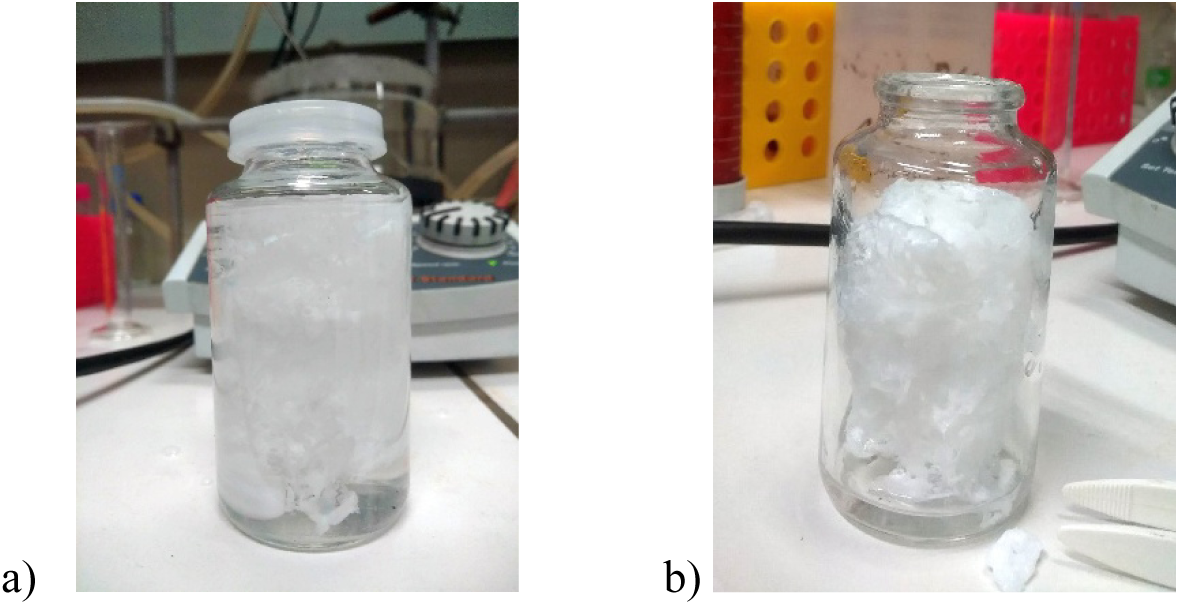
a) Macroscopic precipitate obtained by pouring a PLA solution in THF into hot water ; Fibrous solid obtained after drying overnight, thereafter used as starting material for the cryo-milling process.

**Fig A2:**
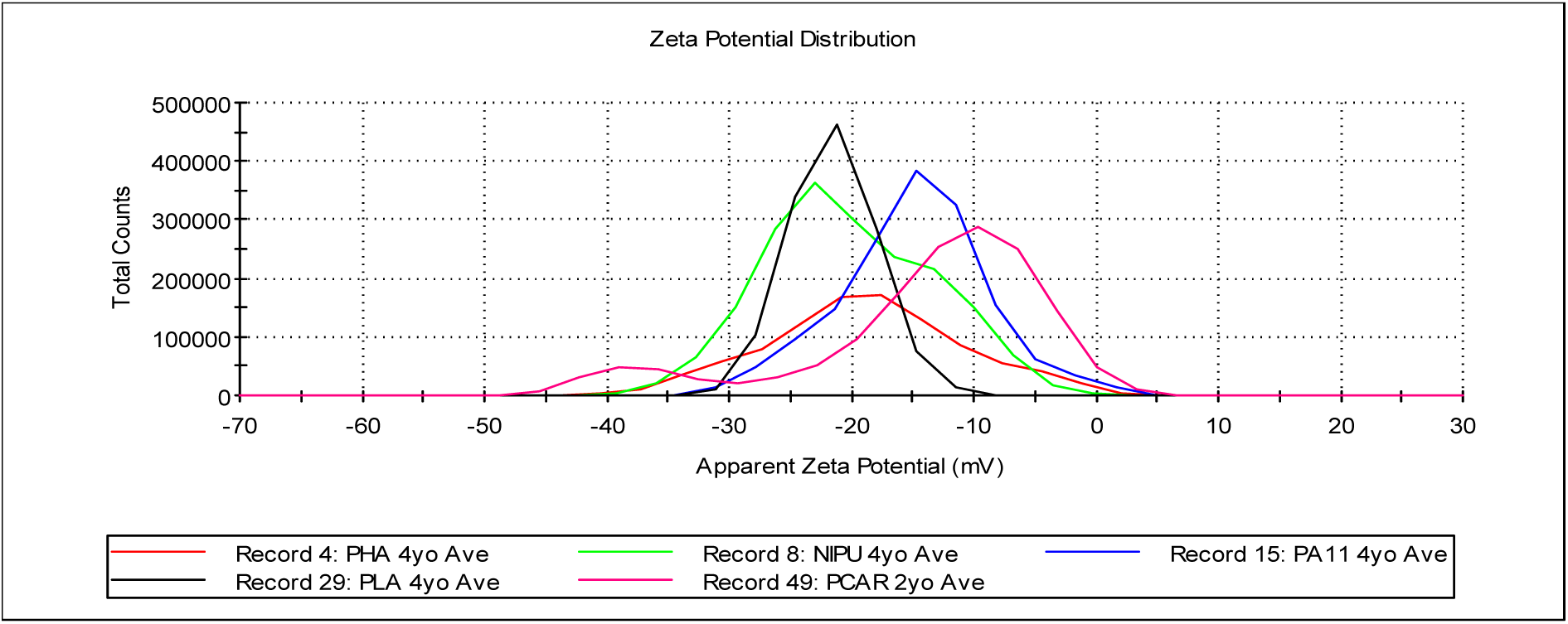
Distribution of Zeta potentials measured by optical phase laser velocimetry at 25°C in water, with a Malvern NanoZS instrument for backscattering DLS and PALS (operating at 173° angle).

## Notes

https://doi.org/10.57745/FKGAPY

### Competing Interest Statement

The authors have declared no competing interest.

### Summary of Updates

This revision contains some text polishing and corrected figures wth their original data made available in an open national repository, in accordance with the policy of the PCI journal to which the work is submitted.

